# Metabolic resistance of Aβ3pE-42, target epitope of the anti-Alzheimer therapeutic antibody, donanemab

**DOI:** 10.1101/2024.01.30.578111

**Authors:** Nobuhisa Iwata, Satoshi Tsubuki, Risa Takamura, Naoto Watamura, Naomasa Kakiya, Ryo Fujioka, Naomi Mihira, Misaki Sekiguchi, Kaori Watanabe-Iwata, Naoko Kamano, Yukio Matsuba, David M.A. Mann, Andrew C. Robinson, Shoko Hashimoto, Hiroki Sasaguri, Takashi Saito, Makoto Higuchi, Takaomi C. Saido

## Abstract

The amyloid β peptide (Aβ) starting with pyroglutamate (pE) at position 3 and ending at position 42 (Aβ3pE-42) is a dominant species that accumulates in the Alzheimer’s disease (AD) brain. Consistently, a therapeutic antibody raised against this species, donanemab, has been shown to be effective in recent clinical trials. While the primary Aβ species produced physiologically is Aβ1-40/42, an explanation for how and why this physiological Aβ is converted to the pathological form has remained elusive. The conversion of Aβ1-42 to Aβ3pE-42 is likely to take place after deposition of Aβ1-42 given that Aβ3pE-42 plaques arise significantly later than Aβ1-42 deposition in the brains of single *App* knock-in and APP-transgenic mice. Here, we present experimental evidence that accounts for the aging-associated Aβ3pE-42 deposition: [1] Aβ3pE-42 is metabolically more stable than other AβX-42 species; [2] Deficiency of neprilysin (NEP), the major Aβ-degrading enzyme, induces a relatively selective deposition of Aβ3pE-42 in APP-Tg mice. [3] Aβ3pE-42 deposition always colocalizes with cored plaques in both APP-Tg and App knock-in mouse brains; [4] Aβ3E-42, an immediate precursor of Aβ3pE-42, as well as Aβ2A-42 and Aβ4F-42 are more short-lived than Aβ1-42 *in vivo*, indicating that simple N-terminal truncation that can arise enzymatically or spontaneously makes AβX-42 easier to catabolize. Consistently, newly generated knock-in mice, *App^NL-(ΔDA)-F^* and *App^NL-(ΔDA)-Q-F^*, showed no detectable Aβ pathology even after aging, indicating that the Aβ3E-42 and Aβ3Q-42 species are extremely labile to the *in vivo* catabolic system and that the E/Q cyclase activity present in mouse brain is insufficient for Aβ3pE-42 generation. In addition, a deficiency of NEP facilitated Aβ3pE-42 deposition. Of note, we identified a trace amount of Aβ3pE-42 and its immediate precursor, Aβ3E-42, in the insoluble fraction of NEP-deficient APP-Tg mouse brains. Aβ3pE-42 is thus likely to be a probabilistic by-product of Aβ1-42 metabolism that selectively accumulates over a long-time range of brain aging. It is likely produced in the solid state or at the solid-liquid interface. Our findings suggest that anti-Aβ therapies will probably be most effective if given before Aβ3pE-42 deposition takes place.

## Text

We previously reported that the majority of amyloid β peptide (Aβ) in the brains of aged humans and a Down’s syndrome patient started with pyroglutamate (pE) at position 3 and ended at position 42 (Aβ3pE-42)(Frost *et al*, 2013; Lemere *et al*, 1996; Saido *et al*, 1995a; Saido *et al*, 1996) (Iwatsubo *et al*, 1996) (Harigaya *et al*, 1995; Kawarabayashi *et al*, 2001). In contrast, transgenic (Tg) mice overexpressing human APP with pathogenic mutations primarily accumulate Aβ1-40 and Aβ1-42 (Kawarabayashi *et al.*, 2001). We demonstrate here, using a panel of antibodies capable of distinguishing among various N-and C-terminal variants of Aβ(Saido *et al*., 1996), that the predominant Aβ species in the brains of Alzheimer’s disease (AD) patients is also Aβ3pE42 (**Supplementary Figure 1**). We estimate that Aβ3pE42 accounts for 50% or more of the total Aβ in AD brains. It is notable that the electrophoretic profiles of AβN3pE and AβC42 reflecting SDS-resistant oligomer formation resemble each other in a specific manner. There is, however, some discrepancy among different reports regarding the quantity of Aβ species, which vary depending on the methods employed. For instance, the group led by Michel Goedert, who used MALDI-TOF mass spectrometry and LC-MS/MS to resolve the Cryo-EM structure of Aβ42 filaments from human brains, recently indicated that Aβ3pE-42 was a minor species in the brains of AD patients and *App* knock-in mice (Yang *et al*, 2022). This inconsistency can be accounted for by the unique physicochemical nature of Aβ3pE-42, which is seldom recovered from reversed phase HPLC under normal conditions (**Supplementary Table 1**). Use of a heated (50 °C) basic solvent containing betaine and limited proteolysis using lysyl-endopeptidase allows full recovery in HPLC and detection by mass spectrometry, respectively. These observations explain the relatively low estimation of Aβ3pE-42 levels in some prior studies as well(Glenner & Wong, 1984, 2012; Mori *et al*, 1992; Wong *et al*, 1985). Consistently, Gunter *et al*.(G✓ntert *et al*, 2006) successfully detected Aβ3pE-42 from human brain by mass spectrometry after proteolytic digestion. It is notable that a therapeutic antibody raised against this species, donanemab, has been shown to be effective in recent clinical trials (Demattos *et al*, 2012; Sims *et al*, 2023).

### Deposition of AβX-42 variants in AD brain

Aβ3pE-42 has been shown to exhibit greater neurotoxicity and oligomerization properties than Aβ1-40 and Aβ1-42 (Dunkelmann *et al*, 2018a; Dunkelmann *et al*, 2018b; Frost *et al*., 2013; Nussbaum *et al*, 2012; Wulff *et al*, 2016). In addition, *App* knock-in mice, which accumulate mainly full-length Aβ1-42, but little Aβ3pE-42, fail to exhibit major subsequent pathologies (tau pathology and neurodegeneration) even after humanization of the entire murine *Mapt* gene(Hashimoto *et al*, 2019) (Saito *et al*, 2019). Generation of this peculiar Aβ species may thus play a major pathogenic role in AD development. The mechanism by which Aβ3pE42 is generated is likely to be mediated by N-terminal truncation of Aβ1-42 by exopeptidase(s), such as aminopeptidases or dipeptidyl peptidase, followed by cyclization of the N-terminal glutamate residue in AβpE-42 (**Supplementary Figure 2**). The cyclization may be spontaneous or enzymatic(Antonyan *et al*, 2018; Cynis *et al*, 2006), but to our knowledge, substantial conversion of Aβ3E-42 to Aβ3pE-42 under physiological conditions has never been demonstrated even *in vitro(Shirotani et al, 2002)*.

Conversion of Aβ1-42 to Aβ3pE-42 results in the loss of one positive and two negative charges at the N-terminus of Aβ(Saido *et al*, 1995b) and may thus account for its unique physical, chemical and biological characteristics. An unresolved problem in understanding the mechanism of Aβ deposition in the human brain, an up-stream event triggering the AD cascade(Selkoe & Hardy, 2016), is the difference in the primary structure of Aβ between the pathologically deposited and physiologically secreted forms(Saido & Iwata, 2006; Saido *et al*., 1995a; Saido *et al*., 1996). For instance, Gravina *et al*. demonstrated using endo-specific antibodies that most Aβ in the AD brain is N-terminally truncated and that Aβ1-42 accounts for only 10-20% of total AβX-42 (Gravina *et al*, 1995), whereas Aβ1-40 and Aβ1-42 are the major species secreted by cells(Scheuner *et al*, 1996; Suzuki *et al*, 1994). The actual amount of the strictly physiological form, Aβ1(L-Asp)-42, in the AD brain is probably even smaller because the N-terminus-specific antibody employed cross-reacts not only with Aβ1(L-Asp)-42 but also with Aβ1(D-Asp)-42, Aβ2A-42 and AβX-42 (X > 3) (Saido *et al*., 1996). **Supplementary Figure 1** shows that, among the structural variants known to be present in AD brains, the Aβ42 bearing amino-terminal pyroglutamate, Aβ3pE-42, is the most abundant in both early- and late-onset cases. This specific form accounts for >50% of total AβX-42 as quantitated against varying amounts of synthetic Aβ peptides, in agreement with Kuo *et al*.(Kuo *et al*, 1997) and Russo *et al.(Russo et al, 1997)*. The presence of Aβ3pE42 in a soluble form even prior to plaque formation in the human brain*(Russo et al., 1997)* indicates the presence of a dynamic equilibrium between the liquid and solid phases.

### Metabolism of isogenic AβX-42 variants *in vivo*

The selective deposition of this physiologically rare Aβ species in human brain can be attributed to its presumed metabolic stability because pyroglutamyl peptide is resistant to major aminopeptidases except for pyroglutamyl aminopeptidase(Mori *et al*., 1992). In accordance with this, an immediate aminopeptidase-sensitive precursor of Aβ3pE-42, Aβ3E-42, is a very minor component in AD brain (**Supplementary Figure 1**). However, our observations of *in vivo* Aβ1-42 catabolism demonstrated that specific endoproteolysis, but not aminopeptidase action, is the major rate-limiting step with Aβ10-37 as a catabolic intermediate(Iwata *et al*, 2000). These observations are better explained by assuming that different Aβ forms have different life spans, for which the structural determinants may reside in the amino-terminal residues of the peptide.

The structural determinants of protein life span have so far been extensively investigated in studies of intracellular proteolysis largely governed by the ubiquitin-proteasome system and autophagy(Kwon & Ciechanover, 2017). The presence of pro-catabolism signals inside substrate proteins such as the “PEST sequence” and “destruction box” as well as the “N-end rule” that correlates the amino-terminal residue to catabolic sensitivity has been described (Dissmeyer *et al*, 2018; Lee *et al*, 2016). However, little is yet known about extracellular protein catabolism, particularly in such complex organs as the brain. This appears to be mainly because the extracellular situations *in vivo* are difficult to reconstitute in a tissue culture paradigm. Thus, elucidating the mechanisms underlying extracellular peptide catabolism profiles as one of the newest frontiers of modern biology and it is of particular interest to see if such rules as the N-end rule do in fact exist.

The present study aims to examine whether the amino-terminal structure of AβX-42 influences its catabolism *in vivo* based on its structural alteration in the AD brain (**Supplementary Figure 1**). The study also serves as an initial attempt to address the question“What determines the life spans of extracellular peptides in the brain?”For this purpose, we synthesized amino-terminal variants of Aβ internally radiolabeled with ^3^H at residues 2, 4, 9, 17 and with ^14^C at 20, 29, 34, 40, 42 and analyzed their catabolic fate in rat hippocampal tissue(Iwata *et al*., 2000). The synthesized variants included Aβ1D(L-Asp)-42, Aβ1rD(D-Asp)-42, Aβ1AcD(Actyl-Asp)-42, Aβ2A-42, Aβ3E-42, Aβ3pE-42, and Aβ4F-42 (**Supplementary Figure 2**). The results shown in **Figure 1** and **Table 1** demonstrate that subtle differences in the amino-terminal structure have a profound influence on the way Aβ is catabolized.

**Figure 1.**
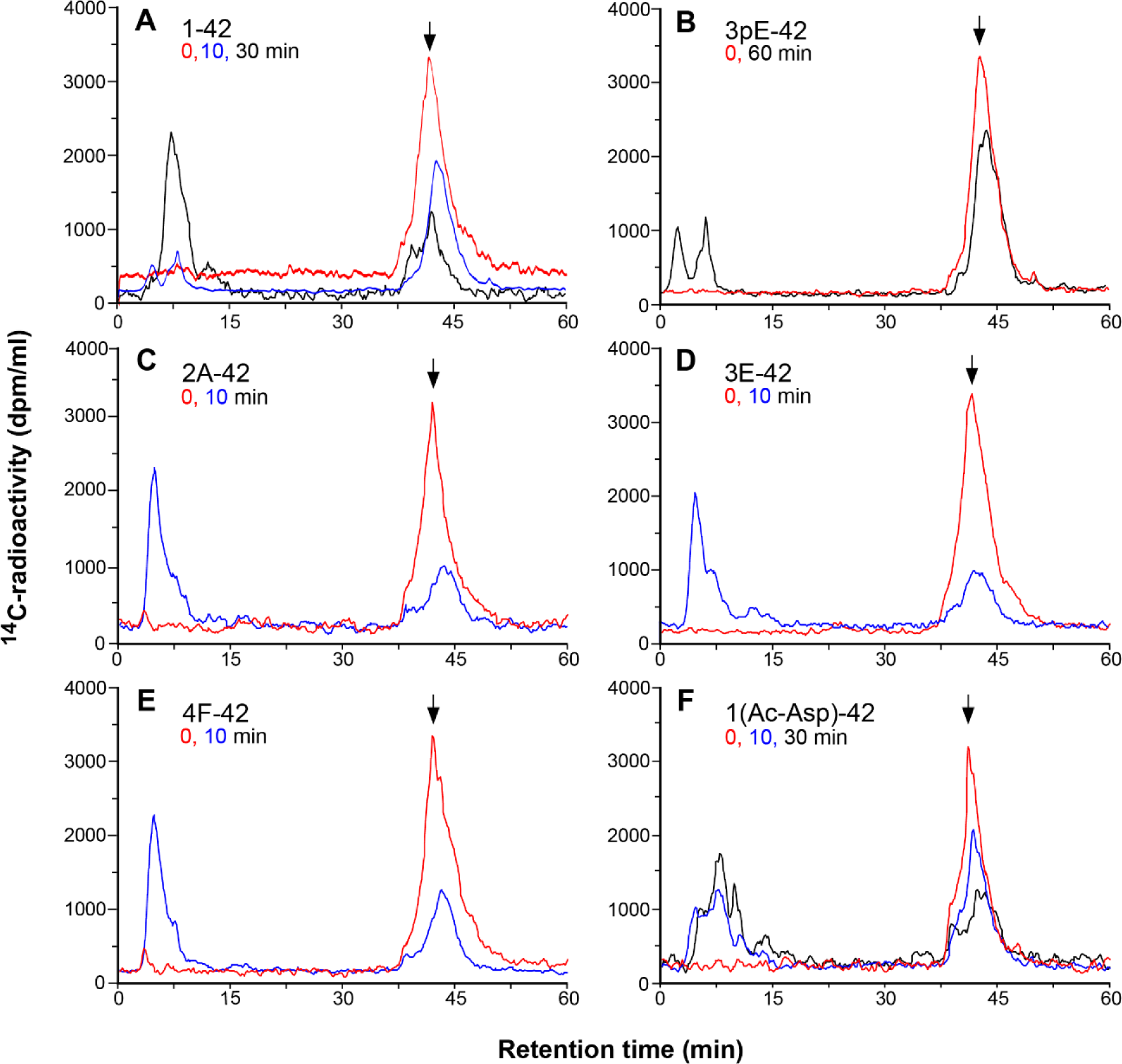
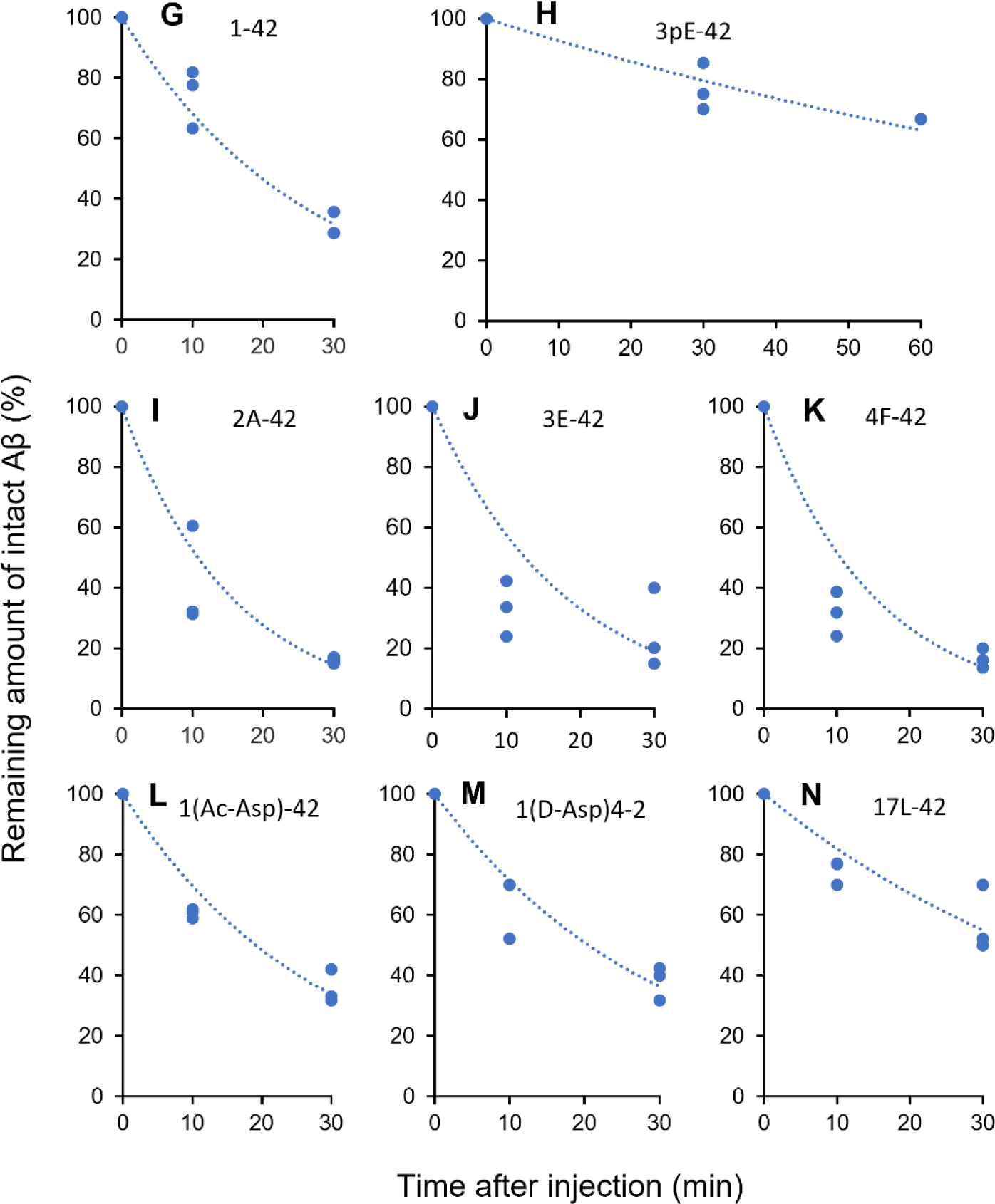
Catabolism of Aβ_X-42_ peptides in hippocampus. *A-F*, Variants of ^3^H/^14^C-Aβ_X-42_ were subjected to *in vivo* degradation and subsequent analysis as previously described(Iwata *et al*., 2000). Briefly, 0.5 μg Aβ_X-42_ peptide dissolved in phosphate-buffered saline was unilaterally injected into the CA1 sector of rat hippocampus and subjected to catabolism until the animals were sacrificed by decapitation at the indicated time points. There was no contamination of the injection area by plasma or cerebrospinal fluid as immunohistochemically confirmed using anti-rat IgG. After extraction, the products were analyzed by the reversed phase HPLC connected to a flow scintillation monitor. The HPLC profiles in ^14^C mode at time 0 (red), 10 (blue) and 30/60 (black) minutes after the *in vivo* injection are shown. The elution profiles in ^14^C and ^3^H modes were essentially identical. The major peaks at time 0 indicated by the arrows correspond the intact substrates. Digitized data were used to calculate the *in vivo* half-lives of the peptides as shown in Table 1. *G-N*, Comparison of in vivo catabolic rates of Aβx-42 variants. Digitized data of remaining amounts of intact Aβx-42 shown in Figure 1 were plotted, and then drawn exponential approximation curves using Microsoft EXCELL.

**Table 1.** *In vivo* half-lives of Aβx-42.

| X (amino-terminal residue) | half-life (min)* | P value against 1(Asp)** |  |
| --- | --- | --- | --- |
| 1(Asp) | 18.08 | Full-length control |  |
| 2(Ala) | 10.77 | 0.02 | significant |
| 3(Glu) | 12.51 | 0.016 | significant |
| 4(Phe) | 10.51 | 0.005 | significant |
| 3(pyroGlu) | 90.25 | 0.005 | significant |
| 17(Leu) | 34.69 | 0.081 |  |
| 1(Ac-Asp) | 19.12 | 0.816 |  |
| 1(D-Asp) | 20.53 | 0.662 |  |

\* Calculated using a formula of exponential approximation curve when the y-intercept was 0.
\*\*Analyzed by two-way ANOVA, followed by *post hoc* test.

First, Aβ3pE-42 was much more resistant to *in vivo* catabolism than Aβ1-42; the majority remained uncatabolized even 60 min after administration into the hippocampus, whereas Aβ1-42 was almost fully degraded within 30 min (**Figure 1, panels A and B**; **Table 1**). This observation is consistent with the presumption attributing the selective deposition of Aβ3pE-42 in human brain to its metabolic stability(Saido *et al*., 1995a). In contrast, Aβ2A-42, Aβ3E-42, and Aβ4F-42 were catabolized even more rapidly than Aβ1-42 **(Figure 1, panels C-E** and **Table 1**). These results clearly indicate the presence of structural determinants of life span in the amino-terminal sequence of Aβ. Again, these truncated Aβ peptides without pyroglutamate are consistently seen as minor components deposited in the senile human brain (**Supplementary Figure 1**)(Saido *et al*., 1996). The catabolic stability of these Aβ peptide species therefore correlates well with their tendencies to be pathologically deposited in the human brain. The only exception is Aβ17L-42 (p3 fragment): It has a longer *in vivo* half-life than Aβ1-42 whereas its quantity in AD brain is small. The reason for this discrepancy is elusive, but we speculate that this particular fragment, lacking the relatively hydrophilic N-terminal half of full-length Aβ, is too hydrophobic to properly analyze in the present experimental paradigm.

These observations also indicate that the loss of the first two amino acid residues, DA, is not the cause of the metabolic stability of Aβ3pE-42. Neither is the loss of the α-amino group because Aβ1AcD-42 was degraded at a rate similar to that of Aβ1-42 (**Figure 1, panel G; Table 1**). The specific structure of Aβ3pE-42 seems to generate an anti-catabolism signal, as discussed later. The consistently facilitated metabolism of Aβ2A-42, Aβ3E-42, and Aβ4F-42 also indicates that the amino-terminal aspartic residue of Aβ1-42 is involved in the regulation of metabolic rate, generating an anti-catabolism signal that is milder than that of Aβ3pE-42. Because the rate-limiting step of Aβ1-42 degradation is catalyzed by thiorphan-sensitive neutral endopeptidase(s)(Iwata *et al*., 2000), the conversion of Aβ1-42 to Aβ3pE-42 is likely to decelerate this process. The facilitation of catabolism by removal of aspartate from Aβ1-42, however, may possibly be accounted for by two distinct mechanisms; one is acceleration of the neutral endopeptidase action and the other is participation of alternative catabolic pathway(s) with respect to Aβ2A-42, Aβ3E-42, and Aβ4F-42, but not for Aβ1-42. It is also notable that the catabolic intermediate identified during the Aβ1-42 degradation (Aβ10-37)(Iwata *et al*., 2000) is missing in the proteolysis of Aβ2-42, Aβ3E-42 and Aβ3pE-42, but not of Aβ1(Ac-Asp)-42 (**Figure 1**). This implies that peptidase(s) other than NEP may participate in the metabolism of Aβ2-42, Aβ3E-42 and Aβ3pE-42.

It is not yet clear why the subtle structural difference between Aβ3pE-42 and AβX-42(X=2A, 3E, 4F) resulted in such a drastic change in the catabolic rate (**Figure1 and Table 1**). One possible factor is an alteration in the tertiary structure caused by increased hydrophobicity: conversion of Aβ1-42 to Aβ3pE-42 results in the loss of one positive and two negative charges among the total of four positive and seven negative charges. In accordance with this, we noted that Aβ3pE-42 is less soluble in water and less recoverable from reversed phase HPLC than Aβ1-42 (**Supplementary Table 1**). He and Barrow also demonstrated that pyroglutamyl Aβ has greater β-sheet forming and aggregation properties(He & Barrow, 1999). The specific hydrophobic nature of Aβ3pE-42 is likely to interfere with the direct or indirect interactions between Aβ and thiorphan-sensitive neutral endopeptidase(s) represented by NEP, which has a catalytic site cavity with a diameter of only approximately 20 angstroms(Moss *et al*, 2018, 2020).

### Effect of NEP deficiency on Aβ1-42 and Aβ3pE-42 deposition in the APP-Tg mouse brain

We subsequently examined the effect of NEP (gene nomenclature: *Mme*) deficiency on N1D and N3pE immunoreactivities in APP-Tg mice (**Figure 2A**). Both N1D and N3pE increased (**Figure 2B**) with a distinct plaque size distribution (**Figure 2C**); the N3pE immunoreactivity tended to be present in smaller/cored plaques. The relative increase of N3pE was significantly larger than that of N1D (**Figure 2D**). It is notable the heterozygous *Mme* deficiency, which corresponds to approximately 50% NEP activity reduction(Iwata *et al*, 2001), resulted in a relative increase in the N3pE/N1D ratio because the NEP expression invariably declines with aging(Hellstrom-Lindahl *et al*, 2008; Iwata *et al*, 2002; Russo *et al*, 2005; Wang *et al*, 2003). We then analyzed the detergent-insoluble/formic acid-soluble fractions of these mouse brains by mass spectrometry (**Figure 3**). Surprisingly, we not only detected N3pE but also a trace quantity of N3E (**Figure 3B**). This observation implies that the Aβ3E-42-to-Aβ3pE-42 conversion may take place in a solid state or solid-liquid interface, which may agree with the following findings. NEP deficiency increased not only the histochemical quantity but also the biochemical quantity of Aβ3pE-42 in the APP-Tg brain (**Figure 4**). When colocalization of Aβ3pE-42 with cored plaques in NEP-deficient APP-Tg mouse brain was examined using Pittsburg Compound B, which known to selectively bind to cored plaques(Ikonomovic *et al*, 2008), (**Figure 5**). PIB binding was increased in NEP-deficient APP-Tg mice in a manner colocalizing with N3pE-positive cored plaques in a statistically significant way (**Figure 5A & B**). Positron emission tomography (PET) analysis also showed an increase of *in vivo* binding of PIB in the NEP-deficient APP-Tg mice (**Figure 5C & D**). These observations consistently indicated that NEP deficiency increases the cored plaque-associated Aβ3pE-42. Incidentally, NEP deficiency also accelerated the deposition of both Aβ1-42 and Aβ3pE-42 in the *App^NL-F^* line of *App* knock-in mice, but there was no significant difference in the ratio of Aβ3pE-42/Aβ1-42. This was presumably due to the presence of the Beyreuther/Iberian mutation in *App* knock-in mice, which increases the ratio of Aβ1-42/Aβ1-40 production(Saito *et al*, 2014) and drives intensive Aβ pathology(Kim *et al*, 2007). In any case, histochemical analysis of APP-Tg (**Figure 2**) and *App* knock-in lines (**Supplementary Figure 4 & 5**) consistently indicated that deposition of Aβ3pE-42 takes place much later than that of Aβ1-42. The biochemical quantity of Aβ3pE-42 accounts for only 0.1% of total Aβ42 in the insoluble fraction of 24-month-old *App* knock-in mouse brains (**Supplementary Figure 4B**).

**Figure 2.**
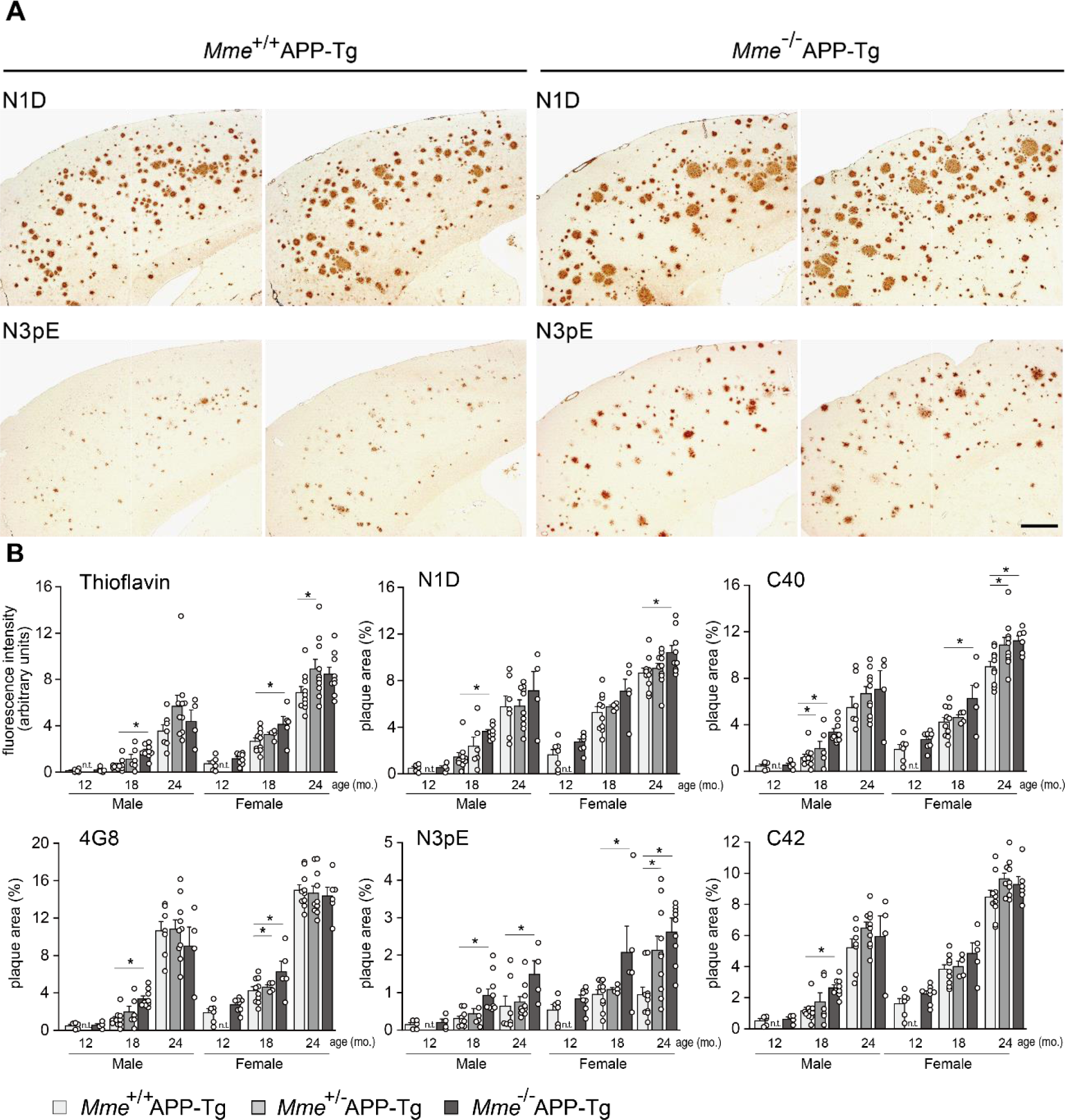
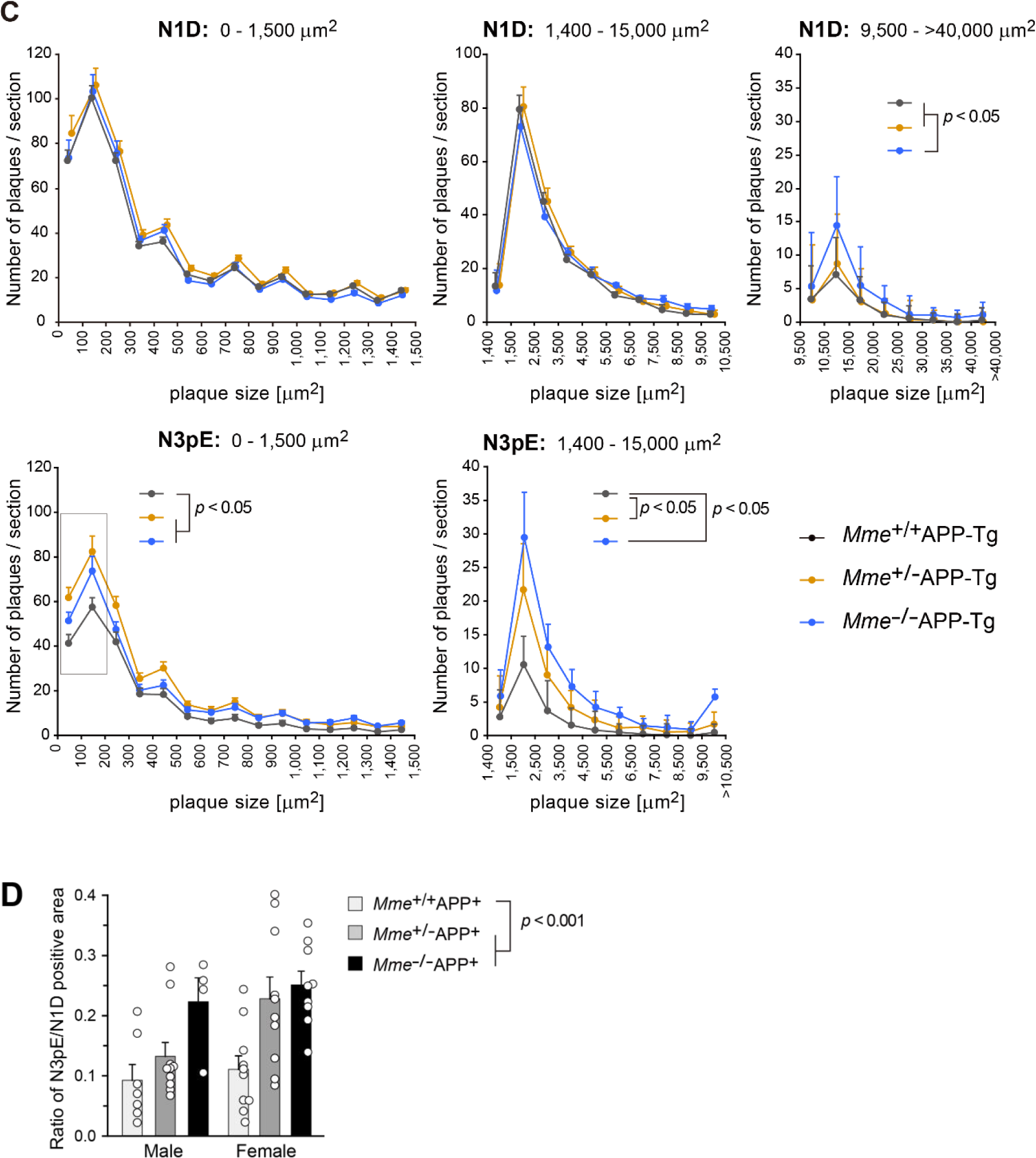
Deposition of Aβ1-42 and Aβ3pE-42 in the brains of aged NEP (*Mme*)-deficient APP transgenic mice. *A,* Immunohistochemical staining of the brains of 24-month-old NEP^+/+^APP-Tg and *Mme*^-/-^APP-Tg mice using anti-Aβ antibodies, N1D and N3pE. Immunostained sections from two individuals in each case are shown. Scale bar, 500 μm. *B*, Aβ deposits in brain sections from *Mme*^+/+^APP-Tg and *Mme*^-/-^APP-Tg mice (12-, 18- and 24-month-old ages) were stained with thioflavin or immunostained with N-terminal specific antibodies for Aβ (N1D and N3pE), anti-pan Aβ antibody (4G8), C-terminal specific antibodies for Aβ (C40 and C42), and then quantified as described in Materials and Methods. Amyloid load is expressed as a fluorescence intensity in the measured area or a percent of measured area. Each column with a bar represents the mean ± S.E. Multiple comparisons were done by a one-way ANOVA, followed by *post*-*hoc* test as described in “Materials and Methods”. Numbers of analyzed animals were as follows: 12-month-old, 5 male and 6 female *Mme*^+/+^ APP-Tg, 4 male and 8 female *Mme*^-/-^APP-Tg;18-month-old, 10 male and 10 female *Mme*^+/+^APP-Tg, 6 male and 4 females *Mme*^+/-^APP-Tg, 9 male and 5 female *Mme*^-/-^APP-Tg; 24-month-old, 7 male and 10 female *Mme*^+/+^APP-Tg, 10 male and 10 female *Mme*^+/-^APP-Tg, 4 male and 9 female NEP^-/-^APP-Tg mice. \**P* < 0.05, significantly different from *Mme*^+/+^APP-Tg mice in the same ages. *C*, Aβ plaque sizes in brain sections from *Mme*^+/+^APP-Tg and *Mme*^-/-^APP-Tg mice (female, aged 24 months) were analyzed using MetaMorph image analysis software. Two or three sections from one individual were analyzed and data were averaged. Area per section analyzed was 13.8 ± 0.058 mm^2^. Each point with a bar represents the mean ± S.E. \**P* < 0.05, significantly different from *Mme*^+/+^APP-Tg mice. Numbers of analyzed animals were as follows: 11 female *Mme*^+/+^APP-Tg, 10 female *Mme*^+/-^APP-Tg; 9 female *Mme*^-/-^APP-Tg. *D*, Ratios of N3pE/N1D-positive areas in *Mme*^+/+^APP-Tg, *Mme*^+/-^APP-Tg and *Mme*^-/-^APP-Tg mouse brains (male and female, 24 months old) were compared. Immunostaining for N3pE and N1D was carried out using 2-3 sets of serial brain sections from each individual, and the average was used as each determinant. Each column with a bar represents the mean ± S.E. The make-up of the animal group (number, males, females, etc) was that same as that shown in Fig. 2B. Two-way ANOVA (gender, genotype) showed a significant effect of the *Mme* genotype (*F*_(2, 44)_ = 9.434; *P <* 0.001).

**Figure 3.**
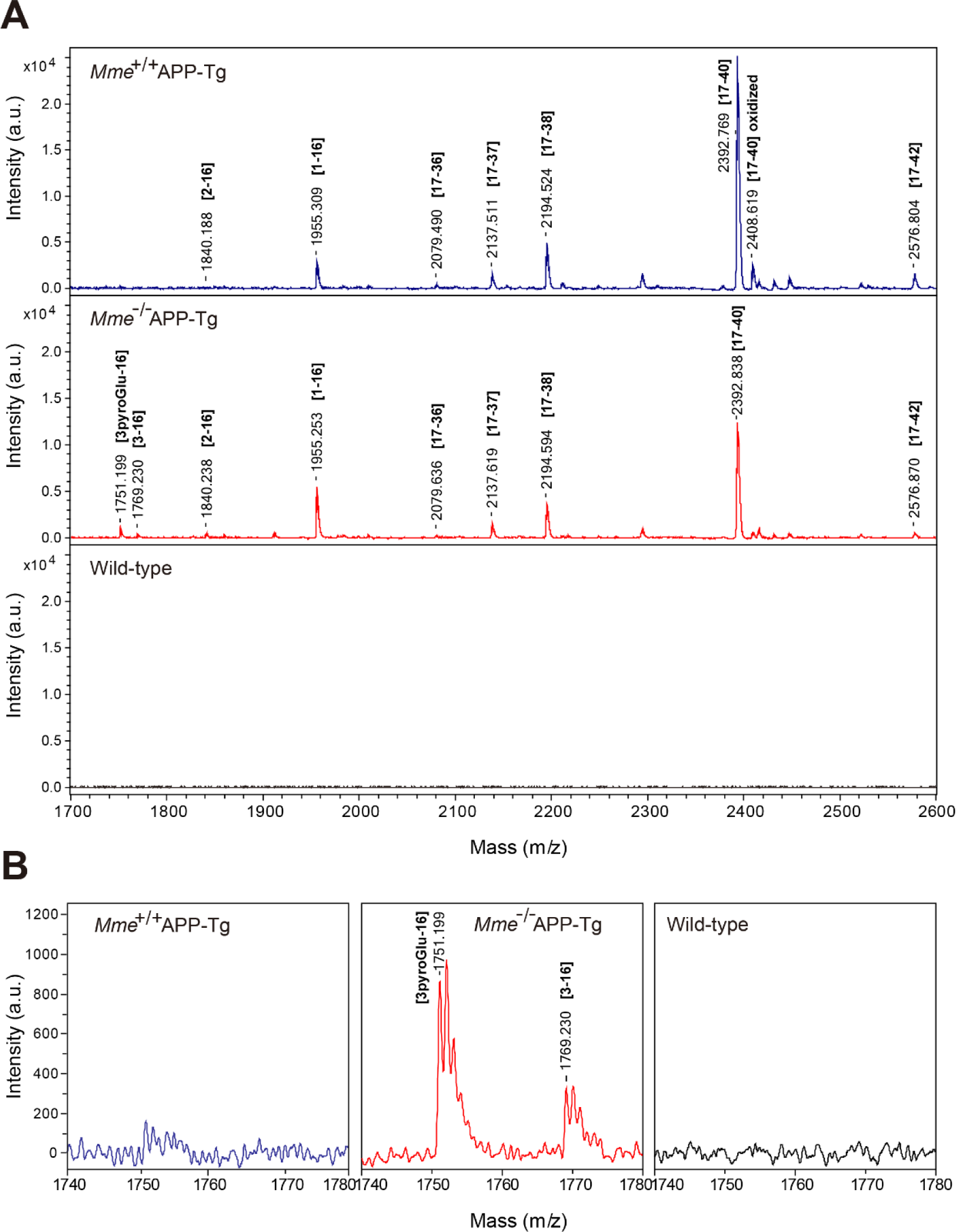
Mass spectrometric profiles from detergent insoluble/formic acid-soluble fractions of brains from aged APP transgenic mice. *A*, Monoisotopic mass of Aβ variants in the detergent-insoluble/formic acid-soluble fractions from 24-month-old APP transgenic and non-transgenic brains were determined after the digestion of lysyl endopeptidase (Aβ with 3pE at the N-terminal shows poor ionization properties). *B*, The m/z range between 1740 and 1780 in Panel A is shown at a higher magnification.

**Figure 4.**
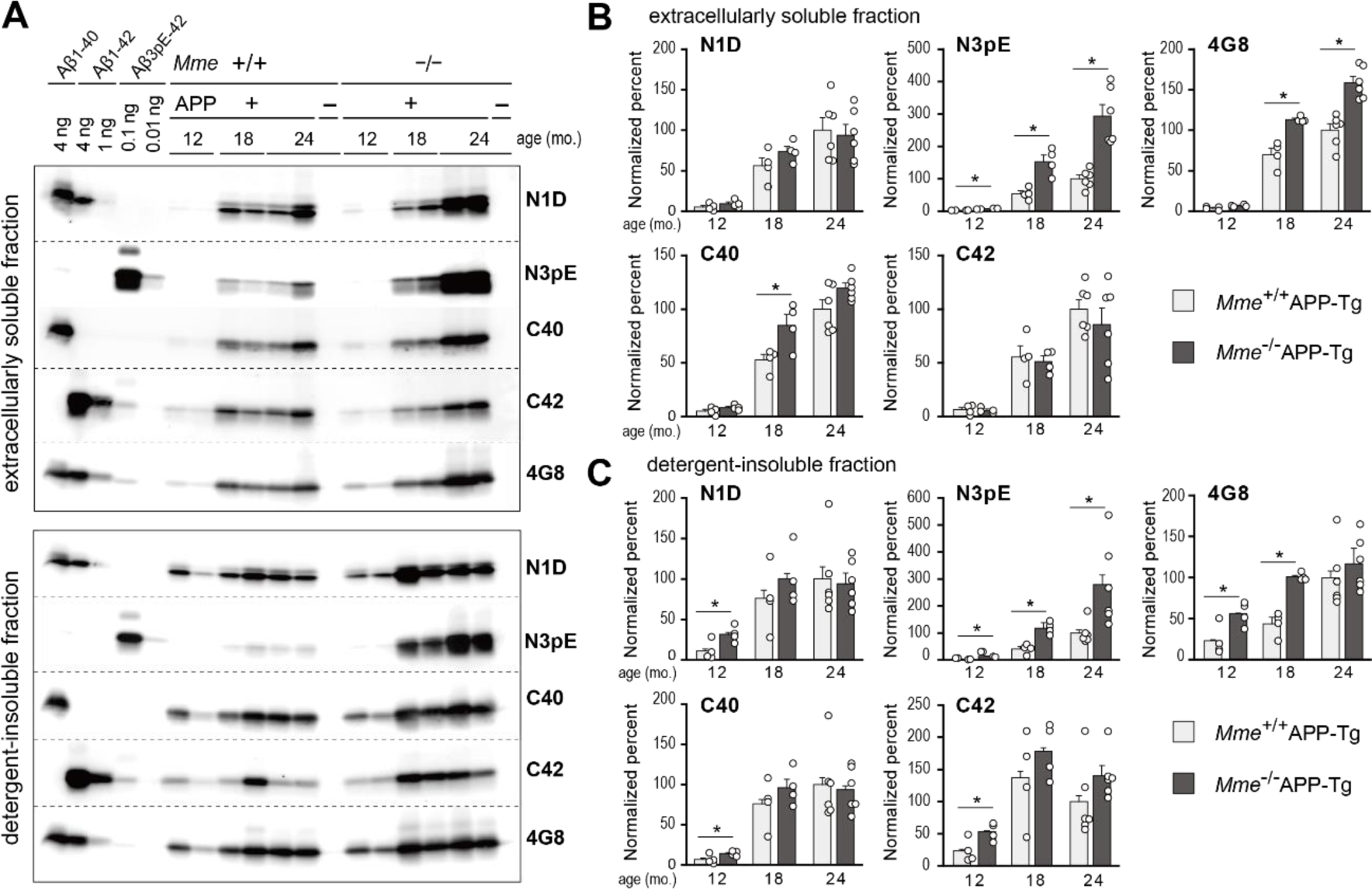
Diverse effects of *Mme* deficiency on extracellularly soluble and insoluble levels of Aβ variants in aged *Mme*-deficient APP-Tg mice. *A*, Brain tissue fractions (upper panel: extracellularly soluble fraction, 0.5 μg protein; lower panel: detergent-insoluble/formic acid-extractable fraction, 20 ng protein) of non-transgenic and APP transgenic mice with or without the neprilysin gene (12-,18-and 24-months old) were subjected to Western blot analyses using N-terminal specific antibodies for Aβ (N1D and N3pE), anti-pan Aβ antibody (4G8), and C-terminal specific antibodies for Aβ (C40 and C42). *B* (extracellularly soluble fraction) and *C* (detergent-insoluble/formic acid-extractable fraction), Intensities of immunoreactive bands on blots shown in Supplemental Figure 2 were quantified as described in Experimental procedures. The intensities were normalized against the data from 24-month-old *Mme*^+/+^APP-Tg mice. The amount of immunoreactive Aβ variant in 24-month-old *Mme*^+/+^APP-Tg mouse brains to N1D, N3pE, 4G8, C40 and C42 is 15.80, 0.028, 10.12, 8.18 and 1.70 ng/μg protein in (*B*), and 241.3, 1.150, 895.7, 461.6 and 91.7 ng/μg protein in (*C*), respectively. Each column with a bar represents the mean ± SEM of 4–6 female mice. \**P* < 0.05, significantly different from *Mme*^+/+^APP-Tg mice at the same ages. Two-way ANOVA revealed that significant interactions between nerilysin-deficiency and ages on N3pE levels in soluble (F_(2,22)_ = 14.612 [*P* < 0.001]) and in the detergent-insoluble/formic acid-extractable (F_(2,22)_ = 1.193 [*P* < 0.05]) fractions.

**Figure 5.**
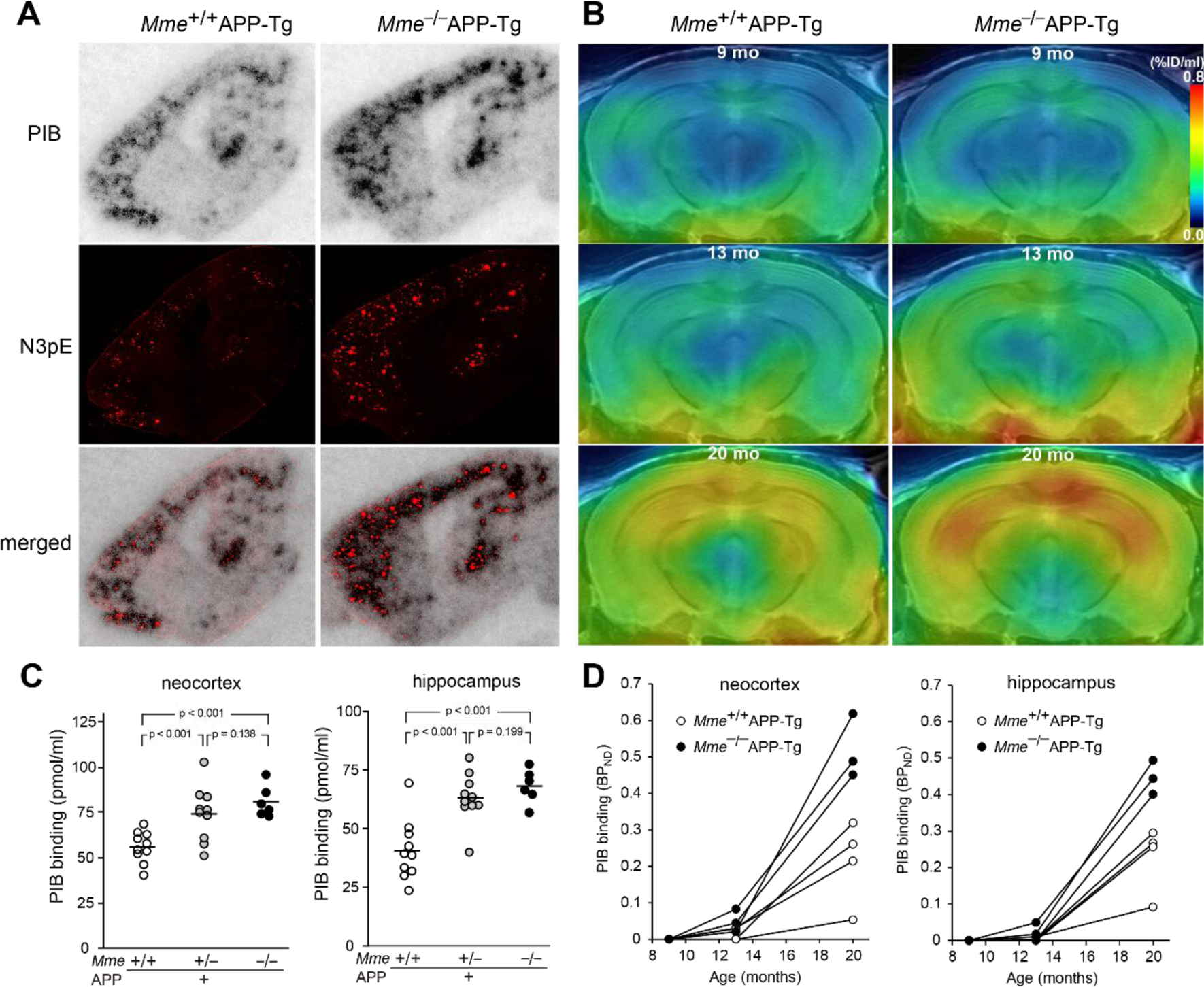
Colocalization of AβN3pE-positive plaques with PIB-binding sites. *A*, Brain sections from 24-month-old *Mme*^+/+^APP-Tg (left) and *Mme*^-/-^APP-Tg (right) mice were subjected to autoradiography with [^11^C]PiB (upper), and thereafter immunostained with polyclonal anti-Aβ3pE antibody (middle). Co-localization of radiolabeling with Aβ3pE deposition is shown in the lower panels. *B*, Intensities of [^11^C]PiB signals (normalized by the intensity of non-specific cerebellar labeling) in the neocortex and hippocampus of 24-month-old APP transgenic mice were significantly elevated and inversely correlated with the gene dose of *Mme*rilysin. Open symbols show individual values obtained by the in vitro autoradiography of brain sections. A solid bar represents the mean value in each group. The following genotypes were tested: *Mme*^+/+^APP-Tg (10 females), *Mme*^+/-^APP-Tg (10 females), and *Mme*^-/-^APP-Tg (6 females). *C*, Time-course PET images of amyloid deposition in the same *Mme*^+/+^APP-Tg (left) and *Mme*^-/-^APP-Tg (right) mice at 9 (top), 13 (middle), and 20 months of age. Coronal images at 3 mm posterior to the bregma were generated by averaging dynamic scan data obtained 30 – 60 min after intravenous administration of [^11^C]PiB, and were coregistered to the MRI template. The radiotracer retention is scaled according to the percentage of injected dose (ID) per tissue volume (%ID/ml). *D*, Longitudinal changes of in vivo [^11^C]PiB retentions estimated as non-displaceable binding potential (BP_ND_) in the neocortex (left) and hippocampus (right) of *Mme*^+/+^APP-Tg (open circles; n = 4) and *Mme*^-/-^APP-Tg (closed circles; n = 3) mice. Data from the same individuals are connected by solid lines. There were significant effects of age [(F(2, 4) = 70.9 and 118.9, p < 0.0001 in the neocortex and hippocampus, respectively) and genotype [F(1, 5) = 18.7, p < 0.01 and F(1, 5) = 15.9, p < 0.05 in the neocortex and hippocampus, respectively] detected by 2-way, repeated-measures ANOVA.

### Generation and analysis of *App^NL-(ΔDA)-F^* and *App^NL-(ΔDA-Q)-F^* knock-in mouse lines

We next aimed to reconstitute Aβ3pE-42 pathology in mouse models, for which two knock-in mouse lines were generated: *App^NL-(ΔDA)-F^* and *App^NL-(ΔDA-Q)-F^* (**Figure 6A**). This strategy was based on our previous experimental results showing that primary neurons expressing APP cDNA with the first two amino acid residues of Aβ deleted (NL(ΔDA)E) produced Aβ3E-40/42 and that those in which E had been replaced by Q (NL(ΔDA)Q) produced Aβ3pE-40/42(Shirotani *et al*., 2002). Indeed, we were able to detect a trace amount (approximately 1 f mol/g) of Aβ3pE-42 only in the brains of *App^NL-(ΔDA-Q)-F^* mice at 2 months of age (**Figure 6**). Despite our expectation, neither of the *App^NL-(ΔDA)-F^* and *App^NL-(ΔDA-Q)-F^* lines exhibited any visible Aβ pathology at all, even after aging (i.e., 18 months or more) (**Figure 6B**). Notably, biochemical quantification indicated that the *App^NL-F^* line deposited more Aβ3pE-42 than the *App^NL-(ΔDA)-F^* and *App^NL-(ΔDA-Q)-F^* lines (**Figure 6C**), consistently indicating that Aβ3E-42 and Aβ3Q-42 are much more short-lived than Aβ1-42 after production *in vivo*.

**Figure 6.**
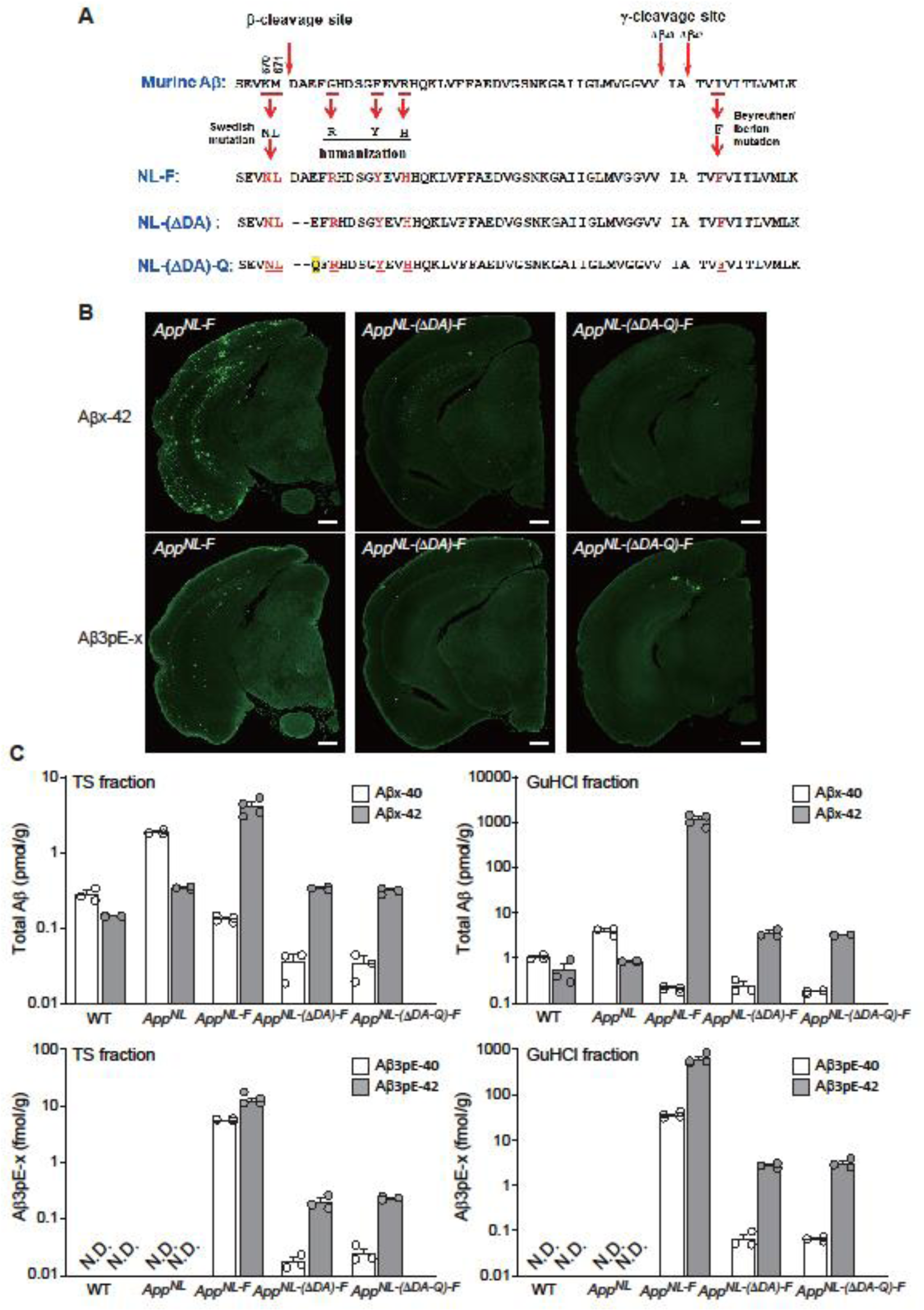
Generation of *App^NL-(ΔDA)-F^* and *App^NL-(ΔDA-Q)-F^* knock-in mice and their Aβ pathology. We generated two additional lines of *App* knock-in mice. In the *App^NL-ΔDA-F^* mice, the first two amino acids of Aβ (DA) are deleted. In the *App^NL-ΔDA-Q-F^* mice, E in *App^NL-ΔDA-F^* is converted to Q. These experimental designs were made based on our previous observation showing that cortical neurons expressing APP cDNAs with the NL-(ΔDA)-Q mutation, but not with NL or NL(ΔDA) mutations, produced Aβ3pE-40/42 (Shirotani *et al*., 2002).

These observations suggest that there may indeed exist an “N-end rule” for Aβ catabolism. The original “N-end rule” for the intracellular protein catabolism is closely associated with a specific class of ubiquitin ligases that recognize the amino-terminal structure of substrate proteins(Dougan *et al*, 2012; Mogk *et al*, 2007; Sherpa *et al*, 2022; Varshavsky, 2017). In the present study, we have demonstrated that the amino-terminal structure of Aβ influences its metabolic rate. Although it is not yet clear how generalizable this “N-end rule” is for extracellular protein catabolism, there may exist a common mechanism that could account for the catabolism of relatively large and hydrophobic peptides represented by Aβ.

## Discussion

Our findings point to the following hypothetical scenario for pathological Aβ3pE-42 deposition in the human brain, as the conversion of Aβ3E-42 to this specific form can be achieved by a single chemical step involving dehydration/cyclization of the amino-terminal glutamate residue (**Supplementary Figure 3**). Under normal conditions, the neutral endopeptidase-dependent pathway, constitutively active, is sufficient to catabolize Aβ1-42 without amino-terminal proteolysis that would produce Aβ2A-42, Aβ3E-42, and Aβ4F-42. This conclusion is arrived at because aminopeptidase inhibitors did not block the *in vivo* degradation of radiolabeled Aβ1-42^(Iwata *et al*., 2000)^ and because Aβ_1(Ac-Asp)-42_ and Aβ_1(D-Asp)-42_, both of which are resistant to mammalian aminopeptidase action (6), underwent *in vivo* degradation in a manner similar to that of Aβ1-42 (**Table 1**). However, under aberrant conditions, presumably caused by a significant reduction of neutral endopeptidase activity or by the presence of an excessive amount of Aβ, a proportion of Aβ1-42 may be processed by aminopeptidases or dipeptidyl peptidases to open the neutral endopeptidase-independent pathway(s), as each class of aminopeptidase is present in the extracellular milieu in brain(Banegas *et al*, 2006; Hui, 2007; Khosla *et al*, 2022) (**Supplementary Figure 3**). Although most of the products would undergo further proteolysis, a small portion of Aβ3E-42 could be converted to the catabolism-resistant form, increasing the probability of pathological deposition. Other possibilities, such as abnormal activation of aminopeptidases, even in the presence of sufficient neutral endopeptidase activity, or aberrant β-secretase activity cleaving at the 2(A)-3(E) site of Aβ in the APP in cells to produce Aβ3pE-42, can also be considered. In any case, the presence of Aβ3pE-42 and Aβ3E-42 in the detergent-insoluble/formic acid-soluble fraction of NEP-deficient APP transgenic mice (**Figure 3B**) implies that the Aβ3E-42-to-Aβ3pE-42 conversion may take place in the solid state or at the solid-liquid interface. Because it is not feasible to spatiotemporally analyze the deposition of Aβ3pE-42 in the human brain, we can only make an assumption based on mouse model data. The relative amount of Aβ3pE-42 per total Aβ in the *App* knock-in mouse model is as small as less than 1% (**Supplementary Figure 4**) and does not recapitulate the massive accumulation of Aβ3pE-42 in human brain (**Supplementary Figure 1**) even if Aβ deposition precedes disease onset by more than 20 years (Bateman *et al*, 2012). Consistently, Yang et al. showed by MALDI TOF mass spectrometry the presence of various Aβ peptides in different AD brains, whereas the *App* knock-in mice uniformly accumulated Aβ1-42(Yang *et al*., 2022), suggesting that Aβ1-42 undergoes truncation over decades after pathological deposition in the human brain. They also found the presence of“Type I and II filaments”in AD brain and only of“Type II”filaments in the *App* knock-in mouse brain. Because both Aβ1-42 and Aβ3pE-42 are abundant in AD brains (**Supplementary Figure 1**) and because Aβ1-42 is the predominant species in *App* knock-in mouse brain (**Supplementary Figure 4**), Types I and II are likely composed of Aβ3pE-42 and Aβ1-42, respectively. Although this assumption needs to be experimentally validated, it may explain why Aβ in *App* knock-in or APP-Tg mice can be extracted by GuHCl(Kawarabayashi *et al*., 2001; Saito *et al*., 2014) whereas that from AD brains requires formic acid (Harigaya *et al*., 1995; Kawarabayashi *et al*., 2001; Saido *et al*., 1995a).

Given that any of these sequential reactions, which are likely to follow a probabilistic-like process, may take much longer than a few years to be pathologically visible in humans, they may be difficult to reconstitute in rodent models, but Aβ3pE-42 appears more stable than other species in both metabolic and structural terms, which is indicative of its selectively low free energy levels. In any case, glutamate cyclization of Aβ3E-42 is a definite final process that results in Aβ3pE-42 production (**Supplementary Figure 3**). Although dehydration/cyclization of glutamate, unlike deamidation-coupled cyclization of glutamine(Abraham & Podell, 1981), has generally been considered to be non-enzymatic, Garden *et al*.(Garden *et al*, 1999) demonstrated the presence of heat-sensitive glutamate cyclase activity in aplysia neurons. If the conversion of Aβ3E-42 to Aβ3pE-42 in human brain happens to be enzymatic, or even to be facilitated by a specific *in vivo* factor, it may become possible to regulate the production of Aβ3pE-42 by pharmacological or dietary means(Coimbra *et al*, 2019; Hennekens *et al*, 2015; Vijayan & Zhang, 2019). Nevertheless, given their nonspecific nature, inhibiting the generation of other physiologically essential pyroglutamyl peptides is likely problematic. Thus, the development of compound(s) that selectively inhibit the conversion of Aβ3E-42 to Aβ3pE-42 based on protein-protein interaction strategies may be imperative. Also, the presence of, for instance, pyroglutamate or derivatives may shift the dynamic equilibrium between Aβ3E-42 and Aβ3pE-42 in a direction that suppresses Aβ3pE-42 formation. Alternatively, the ubiquitously present pyroglutamyl peptidase activity(Abraham & Podell, 1981) would convert Aβ3pE-42 to Aβ4F-42. In any event, it is advantageous that both Aβ3E-42 and Aβ4F-42 are made even more labile to *in vivo* catabolism and less pathologically prone to deposition in the brain than the physiological Aβ1-42 form (**Figure 1 and Table 1**). Also, selective activation of NEP in cortex and hippocampus will be anather therapeutic strategy (See Reference Manuscript 1 and Reference Paper 1).

## Author Contributions

NI, HS, RT, TS, and TCS designed the research plan. NH, HS, RT, NW, NK, RF, NY, MS, KW, YM, and ST performed the experiments. DMAM and ACR collected and provided human brain samples. HS, RT, NK, SH, ST, TS, NI, and TCS analyzed and interpreted data. HS, RT, NW, SH, TS, NI and TCS wrote the manuscript. NI, HS, TO, TS, and TCS supervised the entire research progress.

## Acknowledgements

We thank Taisuke Tomita, University of Tokyo, for valuable discussion. We also thank Yukiko Nagai-Watanabe for secretarial work. This work was supported by research grants from RIKEN, Special Coordination Funds for promoting Science and Technology of STA, CREST, Ministry of Health and Welfare, Ministry of Education, Science and Technology, Chugai Pharmaceutical Co., Mitsubishi Chemical Co., Takeda Chemical Industries, and AMED under Grant Number JP20dm0207001 (Brain Mapping by Integrated Neurotechnologies for Disease Studies (Brain/MINDS)) (TCS) and JSPS KAKENHI Grant Number JP18K07402 (HS).

